# The dispersal of released pheasants and the risk of their intrusion into English protected areas

**DOI:** 10.64898/2026.05.04.722670

**Authors:** Joah R. Madden, Rufus B. Sage, Joe A. Wilde

## Abstract

Large-scale annual releases of pheasants *Phasianus colchicus* and their subsequent management for recreational shooting create various ecological impacts in the UK. While effects at release sites are fairly well understood, dispersing birds may influence areas farther away. If they enter ecologically important but sensitive protected areas (PAs), any negative impacts could be especially harmful. Using tracking data, from 766 birds across 10 sites, we estimated survival and dispersal of released pheasants and applied these patterns to gamebird release records near English PAs to gauge intrusion risk. Of 2,885 registered release sites, just over half lay within 2 km of a PA. A large number of shoots release relatively few birds while a small number release many birds. Thus, numbers expected to enter a particular PA likely depend both on the size of releases and proximity to the PA. We estimate that, at a national level, a maximum of between 525,000 and 784,000 pheasants might be found within PAs very soon after release, representing around 1.7% of all the pheasants released annually. This number declines over the months after release until in February, we estimate that there are between 131,000 and 196,000 pheasants (0.4% of the total release) might be found within PAs. The critical metric by which ecological damage might occur is their density within PAs. Mean densities soon after release averaged 12.0 birds/ha in PAs within 250 m of release sites. This density declined markedly both in time (as birds died) and space (as they moved further from the pen as potential areas increased). By November, densities in PAs 500-1000m from release sites peaked at 0.5 birds/ha, falling to 0.16 birds/ha in February. These estimated densities are around two orders of magnitude lower than those known to cause strong, lasting impacts within release pens. The results are subject to assumptions about movement behaviour, game management and bias in registration. Despite these constraints, considerable local variation exists, with a minority of high-volume release sites very near PAs posing the greatest potential ecological risk.

## INTRODUCTION

In the UK, pheasants (*Phasianus colchicus*), along with red-leg partridges (*Alectoris rufa*) and mallard (*Anas platyrhynchos*), are released in large numbers to provide quarry for recreational hunting (Aebischer 2019, Madden 2021). Typically, they are released into open-topped woodland pens from where they can disperse into the surrounding countryside and return to roost. These pens offer protection from terrestrial predators, primarily red foxes (*Vulpes vulpes*; GCT 1996). The releases occur in July/August when the pheasants, which have been reared under captive condition, are 6-10 weeks old (GCT 1996). The released birds are free to leave the pen at will, but gamekeepers may try to retain them in the area by providing supplementary feed or by “dogging them in” (walking around them to drive them back towards the pen at dusk, often accompanied by dogs). By October, when pheasants can be legally hunted in the UK, the pheasants will have been permitted or encouraged to disperse to outlying areas such as other woodlands or cover crops from where they can be driven back to the pen wood during a hunting day.

Released birds may directly cause negative ecological effects to the fauna and flora in and around release sites. The pheasants can increase soil and water nutrient levels, directly damage flora, predate local species, compete for resources with non-game species, and provide a source or harbour for pathogens and parasites that may infect other species (reviewed in Madden & Sage 2020, Mason et al. 2020, Sage et al. 2020, Madden 2023a). These negative effects are likely to be stronger at higher densities of released birds (e.g. Sage et al. 2005a, Gortazar et al. 2006, Pressland 2009, Neumann et al. 2015, Capstick et al. 2019, but see Davey 2008). Changes in soil chemistry and plant species richness can persist at woodland release sites for around a decade after releases cease, but this may be longer if the densities of released birds are >1000 birds/ha (Capstick et al. 2019). Evidence for these effects has so far focused on densities within the release pens themselves.

The direct negative effects of released gamebirds may be of particular concern if they occur in habitats that contain ecologically important features that are susceptible to damage from gamebirds. Such habitats may be under a range of legal protections. In the UK, some of the most important sites include Special Areas of Conservation (SAC, designated under the Habitats Directive) and Special Protection Areas (SPA, designated under the Birds Directive). Collectively, these form part of the Natura 2000 network that spans Europe. The UK government is under obligation to prevent deterioration of these habitats as well as significant disturbance of the designated species in Natura 2000 sites. A third set of sites, Sites of Special Scientific Interest (SSSI), may also be subject to negative direct effects if they contain designated features susceptible to gamebird activity. In England, there is a recognition of the potential effects of gamebirds being released in or near these protected areas and legislation is in place to attempt to reduce potential risks. Releases inside, or within a buffer zone of 500 metres, of Natura 2000 sites currently require a licence. This may be a general licence (GL43 for SACs, GL45 for SPAs covering pheasants and partridges) if a series of conditions relating to release densities (<700 birds/ha of release pen/area within the site or <1000 birds/ha of release pen/area within the buffer zone), release site habitats, feeding sites and reporting requirements can be met. If these conditions cannot be met, then an individual licence must be applied for. (DEFRA, 2025). Releases and aspects of their subsequent management in SSSIs may need consent from Natural England (DEFRA, 2024). This legislation is designed to reduce densities of released gamebirds within protected areas. However, releases outside protected areas or their buffer zones are not subject to licence or consent, but birds dispersing from them could intrude into protected areas and cause negative ecological effects. The magnitude of these effects will likely depend on the density of birds that occupy the protected area.

Stakeholders (including nature reserve staff, farmers and gamekeepers) hold a variety of opinions on what the effects of release and management might be on protected areas. However, they broadly agree that there are benefits of hedgerow creation and management accompanied by negative effects arising from increased generalist predator populations (Minter et al. 2024). Site-specific case studies suggest that high-density releases in or close to protected areas may alter aerial nitrogen levels, impacting on lichens (Bosanquet 2019), adder (*Vipera berus*) populations (Hand 2020), or change soil nutrient composition and reduce bryophyte abundance (Rothero 2006). We expect that any such effects will scale with release numbers and densities in protected areas.

Although the density of birds at release can be calculated accurately, by considering the numbers of individuals and size of the area or pen where they are placed, accurate density measures become harder to determine as mortality and movement soon mean that numbers of living birds decrease and birds disperse from the area, meaning densities change in the release area and surrounding landscape. Released gamebirds suffer relatively high mortality soon after release, typically in July/August and during the shooting season (finishing at the end of January), although mortality decreases as the year progresses (Madden et al. 2018, Sage et al. 2018). Mortality may occur due to predators, disease or hunting. At the start of the breeding season in the year after release, approximately 9-15% of total released pheasants are still alive (Madden et al. 2018), with <5% of the released birds remaining alive a full year after release (Sage et al. 2001). Therefore, the number (and hence density) of birds is expected to decline markedly over the months after release due to natural mortality or shooting. In addition to mortality, the density of birds in an area is affected by their movement behaviour. Pheasants in the UK exhibit preferences for particular habitats both during the winter (Hill & Ridley 1987, Robertson et al. 1993a) and in the breeding season (Lachlan & Bray 1976, Robertson et al. 1993b). Pheasants are predominantly terrestrial and frequently move along linear features such as hedges, ditches, woodland edges or watercourses (Hill and Robertson 1988, Beardsworth et al. 2021). Thus, their dispersal may not be random in space. With age, they become more susceptible to density-dependent social pressures that drive them from the pen (Whiteside et al. 2019). In addition, gamekeepers may deliberately try to encourage or prevent birds from moving to particular areas (GCT 1996). Pheasants initially stay close to their release site, either because it offers them security and resources, or because gamekeepers deliberately try to constrain them to the area. These factors are likely to vary markedly between release sites, individual keeper activity and time. However, as they are eventually encouraged to disperse, their density inevitably decreases as a square of the distance from the release pen.

Only one study that we are aware of has attempted to assess the densities of pheasants outside release pens. Sage *et al*. (2025) investigated ten sites in Southern England managed for shooting, finding median pheasant densities of around 500/km^2^ post- release (max density approaching 2500/km^2^), falling to around 450/km^2^ during the shooting season, with typical densities falling to 100-250/km^2^ in the same areas during the Spring when the shooting season had ended, and 30-60/km^2^ in the summer. By contrast, the densities of pheasants 2km away from release sites (recorded at eight locations) were typically <20/km^2^ at time of release, rising to 20-45/km^2^ during and at the end of the shooting season, and declining to 5-20/km^2^ in the summer.

It is important to understand what sorts of densities we might expect to find in protected areas in England, and calculating these densities depends on three factors. First, we require the location of release sites relative to the protected areas. In the UK, all gamebird keepers have to register the location of their holdings, the number and species of birds that they hold, and the purpose for which the birds are held, in the Poultry Register. This provides us with a set of locations where gamebirds are released. Second, we require the number of birds being released at a location. These data can also be obtained from the Poultry Register, by considering the records for the numbers of pheasants released, or by considering the records for the numbers of all pheasants held for any purpose. Third, we require a measure of how far released birds are likely to move, how long they might survive and how long they may occupy an area. By tracking a series of cohorts of released pheasants over a range of sites and years, we can determine the proportion of released birds likely to be alive each month and where they have moved to in that time. While precise movement patterns depend on local habitat, environmental and climatic conditions, and game-management actions, which we cannot model at a national level, we can assume general dispersal patterns. Therefore, we can make a series of calculations to estimate how many released pheasants might be found in protected areas in England at different times of the year by considering (i) where and how many birds are released, (ii) how far the birds might disperse, given movement and survival, and (iii) how far these release sites are from protected areas.

## METHODS

### i. Location and Size of Pheasant Releases in England

In the UK, all gamebird keepers have to register the location of their holdings, the number and species of birds that they hold, and the purpose for which the birds are held, in the Poultry Register (APHA & DEFRA 2018). This provides us with a set of locations where gamebirds are released. A weakness of this dataset is that typically only a single location is given per site, whereas shoots typically release their birds at multiple pens around a central location. Consequently, the precision of the location data from the Poultry Register is likely to be low, with potential error of several hundred metres. Another weakness is that, despite being a legal requirement, there is low compliance with or inaccurate reporting to the Poultry Register, with estimated compliance at 28%-73% (Madden 2023b). The total numbers of pheasants recorded as being held for release in the UK in 2024 was 12.2 million, which is only 39% of the 31.5 million pheasants estimated to be released by Madden (2021) or 26% of the 47 million pheasants estimated to be released by Aebischer (2019). Consequently, the accuracy of the release number data from the Poultry Register is likely to be low. Therefore, we extrapolate to national numbers by correcting the numbers reported in the Poultry Register by estimates by Aebischer (2019) and Madden (2021).

We took locations of release sites and release numbers from the Poultry Register data on 7 February 2025 via an enhanced Data Sharing Agreement. We filtered the data so that we could use records from sites that reported holding pheasant, for releasing, for shooting or other purposes (Livestock Unit Animal Production Usage = “Shooting” or “Other”; Livestock Unit Animal Purpose = “Releasing for Shooting”). We concentrated on release sites in England. This dataset included 2,885 locations where pheasants were reported as being held for release, totalling 9.96 million birds.

### ii. Determining the Proximity of Protected Areas to Release Sites

We used published GIS layers depicting protected areas in England/Great Britain. These comprised: GB Special Areas for Conservation SAC (excludes offshore areas, JNCC 2025); GB Special Protect Areas SPA (includes offshore areas, JNCC 2024) and English Sites of Special Scientific Interest SSSI (Natural England, 2025). There is some overlap between these sets of protected areas. All SAC are also SSSI, but not all SSSI are SAC. Some SPA are SSSI, but others are not. We masked these to include only sites in England using a UK boundary map (Igismap, 2025). We calculated how many hectares of protected areas sat within each of the 4 distance bands (250m, 500m, 1000m, 2000m) from a release site, for each of the 2885 reported release sites. From this, we could calculate the proportion of each distance band that was comprised of protected area of each type.

### iii. Pheasant Survival and Dispersal

We used tracking data from 766 pheasants, released at 10 sites in six counties (Table 1), from three other studies. Birds were tracked using a range of technologies, for up to eight months (July-February) between 2001 and 2024, to document month-by-month patterns of mortality and distances dispersed from the release site.

**Table 1.** Numbers of pheasants tracked in each year in each cohort at sites participating in the three studies.

| Study/Site | Year of release | Number of pheasants tracked | Median location frequency |
| --- | --- | --- | --- |
| Mid Devon site | 2017 | 124 | 5 minutes |
| Mid Devon site | 2018 | 126 | 5 minutes |
| Yorkshire site 1 | 2023 | 10 | 60 minutes |
| Yorkshire site 2 | 2023 | 10 | 60 minutes |
| Yorkshire site 3 | 2023 | 10 | 60 minutes |
| Berkshire site (f) | 2001 | 27 | 2 days |
| Hampshire site 1 (b) | 2001 | 24 | 2 days |
| Wiltshire site (cf) | 2001 | 30 | 2.1 days |
| Hampshire site 2 (cr) | 2001 | 30 | 2.1 days |
| Dorset site (h) | 2001 | 29 | 2.8 days |
| Hampshire site 3 (w) | 2001 | 25 | 2 days |
| Berkshire site (f) | 2002 | 24 | 2.1 days |
| Hampshire site 1 (b) | 2002 | 25 | 2.1 days |
| Wiltshire site (cf) | 2002 | 28 | 2.4 days |
| Hampshire site 2 (cr) | 2002 | 29 | 2.4 days |
| Dorset site (h) | 2002 | 30 | 2.1 days |
| Hampshire site 3 (w) | 2002 | 27 | 2.1 days |
| Berkshire site (f) | 2003 | 24 | 2.1 days |
| Hampshire site 1 (b) | 2003 | 25 | 2.2 days |
| Wiltshire site (cf) | 2003 | 29 | 2.1 days |
| Hampshire site 2 (cr) | 2003 | 27 | 4.7 days |
| Dorset site (h) | 2003 | 28 | 2.9 days |
| Hampshire site 3 (w) | 2003 | 25 | 2.1 days |

**Table 2.**
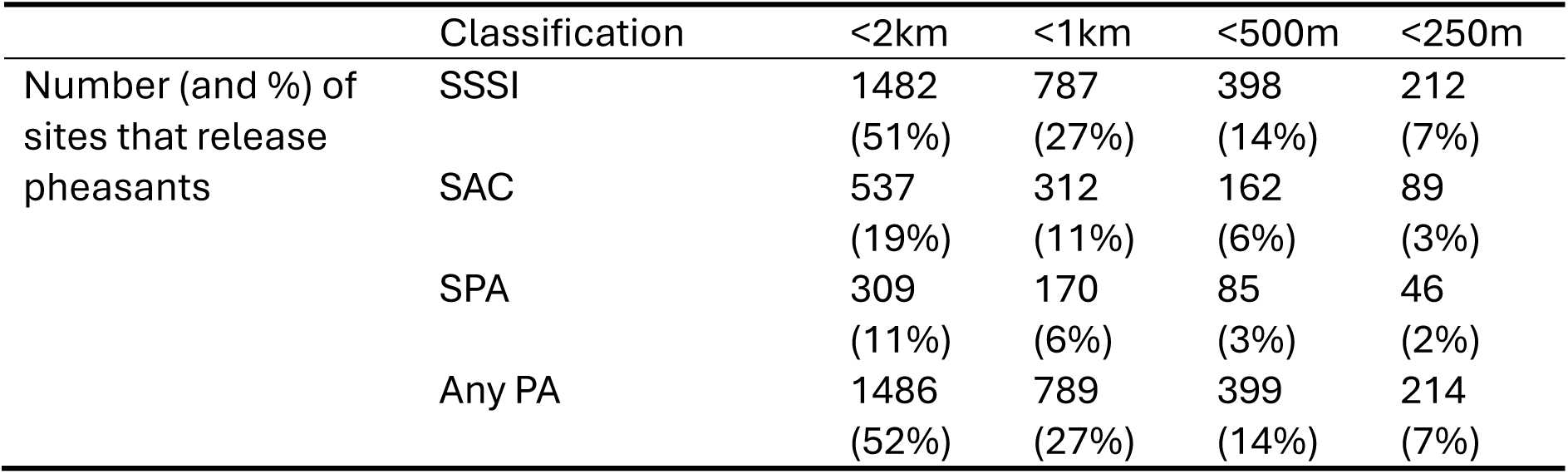
The numbers of the 2885 sites reporting holding gamebirds for release that are located at different distances from English Protected Areas (SAC, SPA and SSSI).

First, 250 pheasants (123 male, 127 female) were tracked at a single site in mid Devon at high spatial and temporal resolution using a reverse GPS tracking system, ATLAS, described and validated in Beardsworth et al. (2022). The tracking system was installed for a series of other studies (e.g. Beardsworth et al. 2021, Heathcote et al. 2023) and data was collected from the site over two seasons 2017-18 and 2018-19. Locations were recorded remotely at 1/4Hz with pooling of locations to improve spatial accuracy at the expense of temporal resolution such that location fixes were obtained every five minutes. Birds were fitted with a backpack tag (22g) and released when ∼9-10 weeks old. The site was a research farm where birds were released from a single, central woodland pen from where they could disperse freely into an area containing 43 artificial feeders. For the first two months post-release, birds were driven back towards the release pen at dusk. There was no predator control on the site nor any game shooting. However, shooting of game and predators occurred on neighbouring farms within 2km of the release site.

Second, 30 pheasants were tracked at three sites in northern England using Lotek GPS tags (PinPoint GPS Argos tags) as part of a study by the Animal and Plant Health Agency specifically to explore the movement and survival of released pheasants. The study was conducted from release in 2023 until March 2024. At each site, 10 birds were fitted with a backpack tag (18g) and released from a single woodland pen when 7-11 weeks old, according to estate management practice. Location fixes were recorded each hour during both daylight and the night. The sites were operating game shoots that had agreed to participate in the study and where a variety of gamekeeping management techniques were used including artificial feeding, dogging in and predator control.

Third, 486 pheasants were tracked at six professionally managed game shoots in southern England using VHF radio tags that were located manually as part of a study into the management of released gamebirds (Turner 2007). The study was conducted over two years with a new cohort of birds released when ∼7 weeks old and tracked before and during the game shooting season. Attempts were made to locate each bird at least twice a week, but other locations were also collected where possible. This gave us mortality data precise to ∼3 days and movement data for all tracked birds. The sites were operating game shoots that had agreed to participate in the study and where a variety of gamekeeping management techniques were used including artificial feeding, dogging in and predator control.

Determining mortality from tracking data is not straightforward (Sergio *et al*. 2019). In some cases, carcasses could be located, either via searches at the last known location (ATLAS and GPS systems) or searches for VHF tags equipped with mortality switches when these were activated. However, tags were sometimes found separate from the bird, and it was difficult to confirm whether they had been shed by a live bird or had been torn from a dead bird by a predator and discarded. In other cases, tags were assumed to have malfunctioned, and the bird being tracked was lost. The probabilities of each of these fates occurring likely differed between the datasets and between sites. Therefore, we took the last day on which a definite signal of life (movement between locations on consecutive fixes) was recorded as the day of death. Given the high temporal resolution of the ATLAS and GPS system, we are confident that our assigned mortality days are accurate to within 24 hours. For the VHF tracking data, average intervals between fixes were 2.35 days and thus our precision of mortality is likely to be accurate to the nearest 2-3 days.

To determine the spatial distribution and dispersal of tracked pheasants, we divided the landscape surrounding each of the release pens into 4 concentric distance bands: 0 – 250m, 250 – 500m, 500 – 1000m, and 1000m – 2000+m and evaluated the number of fixes recorded in each band per month. Because the location fix rate varies across each data set (and between individual tags), and the number of birds alive declined with time due to mortality, we converted the number of location fixes in each band to a proportion of the total number of fixes recorded in that month at each site. This provides an index of potential ecological impact, corresponding to the occupancy density of birds in an area: The more birds in an area and the longer that they spend there, the higher the risk of direct negative ecological impacts. We processed the fix data in four steps. Step one: we calculated for each month, and for each of the three data sets separately, the proportion of all the total fixes for that month and study that were recorded in each of the four distance bands. Step two: for each study, we then overlaid the mortality profile, multiplying the proportion of monthly fixes by the proportion of the original birds still alive in that month. This accounted for the differential fix rates and mortality patterns observed on each study. Step three: we then combined data from the three studies, weighting the values calculated in step 2 by the number of birds contributing data in each study. Therefore, in August, the total of the proportion of these mortality and sample size corrected fixes across the four distance bands summed to 1, while in February, because birds had died, the total was 0.166, reflecting a decrease in number of birds.

We used these proportions as a gross, nation-wide representation of dispersal patterns to calculate the total number of birds likely to be found at different distances from the 2885 release sites that reported releasing pheasants in the Poultry Register. We did this by simply multiplying the original reported number of birds released by the proportions for each month in each distance band. From these values we could estimate the number of released pheasant that might be found in the protected areas within 2km of each release site. We did this by multiplying the number of birds found in each distance band in each month by the percentage of area of the distance band that comprised of protected area (separating out SSSI, SPA and SACs) as calculated and described in §ii).

We then attempted to scale these estimates to a national level by accounting for underreporting of release numbers. At the 2885 sites we included, 9.96 million pheasants were reported as being released so we accounted for the differences to the numbers estimated as being actually released (see §1) by multiplying vales by 2.58 if the Madden (2021) estimate is applied or 3.85 if the Aebischer (2019) estimate is applied.

Direct ecological impacts are strongly related to the density of birds, but as a given number of birds disperse further from the release point, the total area that they might occupy increases and thus the density of birds in that area decreases. To determine density of released pheasants occurring in each protected area in each month, we took the estimated number of pheasants found in the distance band (as described above) and divided it by the area (in ha) of that band that comprised Protected Area (separating out SSSI, SPA and SACs) to give density measures in birds/ha that could then be compared with density levels previously reported to provoke direct negative ecological effects. We focussed on densities based on current best-practice values of 700 birds/ha and two orders of magnitude less than this (70 & 7 birds/ha) to allow the reader to scale likely effects.

All calculations were conducted in R 4.4.1 (R Core Team 2022) using the sf package 1.0-19 (Pebesma & Bivand 2023).

## RESULTS

### i. Location and Size of Pheasant Releases in England and their Proximity to Protected Areas

There was a strong skew in the distribution of the sizes of pheasant releases across England (Figure 1). A large number of shoots release relatively few birds while a small number release many birds. Of the 2885 sites that reported holding pheasants for release, 1638 (57%) reported holding one thousand or fewer. Half (∼4.98 million) of the total number (∼9.96 million) of birds reported as being held for release were located at 158 sites, comprising just 5.4% of the total registered sites. The highest number of pheasants reported as being held for release at a single site was 150,000.

**Figure 1.**
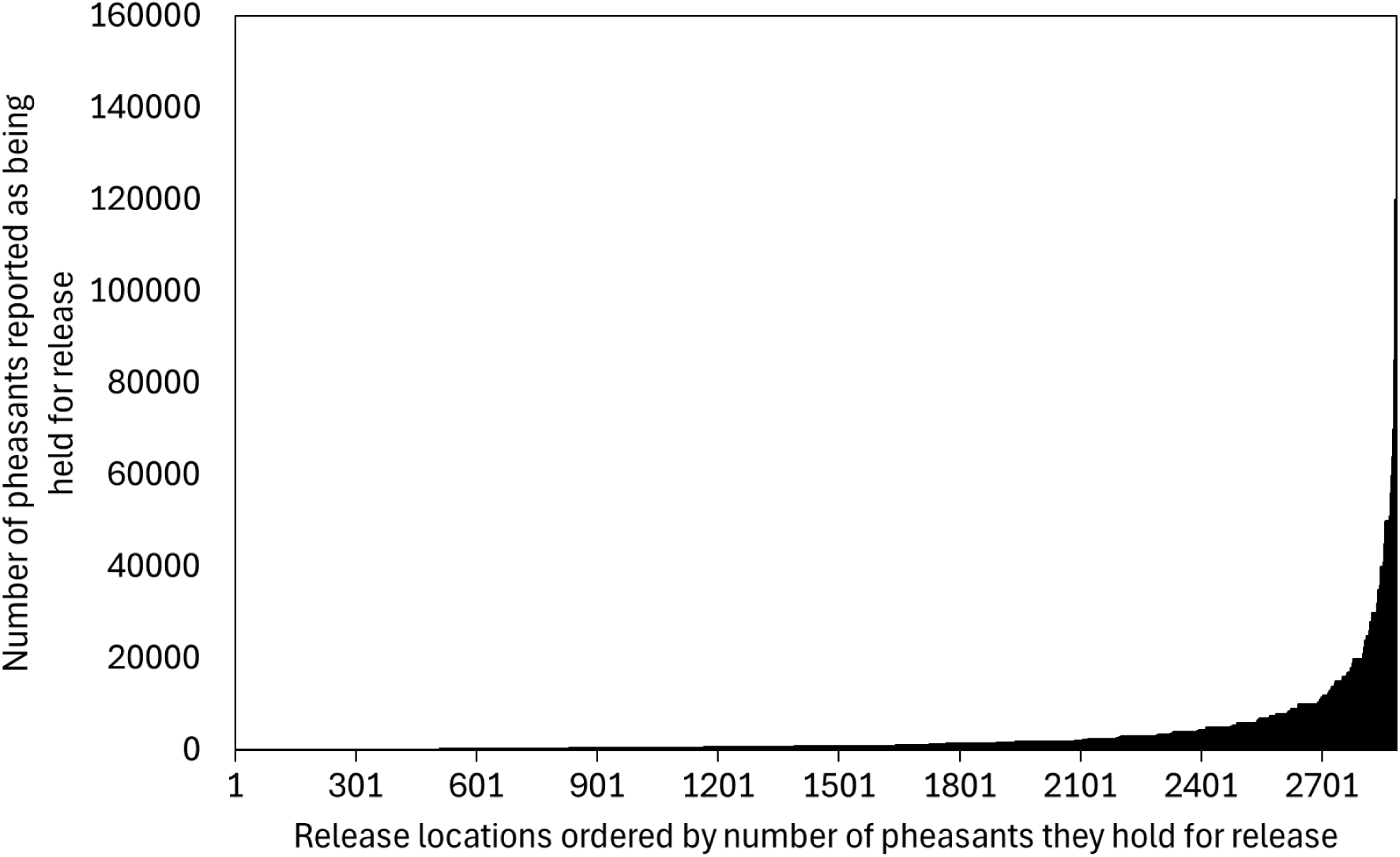
The distribution of the numbers of pheasants reported as being held for release at 2885 sites in England in in the 2024 Poultry Register. The sites are arranged along the x axis in order of their reported size of releases from the smallest releases on the LH side to the largest releases on the RH side.

Of the 2885 sites where pheasants were reported as being held for release, 14% were less than 500m from a SSSI, 6% were less than 500m from an SAC and 3% were less than 500m from an SPA, and 14% were less than 500m away from any PA (Table 1). 52% were within 2km of any protected area. On average, sites where pheasants were recorded as being released had 1.4ha (1SD = 6.2ha) of protected area (SSSI) within 500m, including 0.5ha (3.7ha) of SAC and 0.5Ha (4.6ha) of SPA. Within 2km of release sites there was a mean of 41.2ha (106.8ha) of protected area (SSSI), including 20.3ha (78.2ha) of SAC and 20.1ha (90.2ha) of SPA. The total protected area within 2km surrounding the 2885 sites reporting holding pheasants for release area comprised 118546ha (SSSI) including 58517ha of SAC and 57870ha of SPA. We estimate that these 2885 sites represent around one third of the 7000-9000 shoots in the UK.

### ii. Pheasant Dispersal and Intrusion into Protected Areas

The records of released pheasants at different distances changed markedly throughout the season due to dispersal and death. In the first month after release (August), 95% of monthly location fixes from the tracking data sets (weighting for numbers of birds tracked) were within 500m of the release pen (Figure 2). By November, birds had started to disperse away from the pen and been subject to at least one month of hunting, such that with 44% of the original numberl of fixes (weighting for numbers of birds tracked and accounting for deaths over time) from that month were recorded within 500m of the pen, and 26% of the original level of fixes were recorded further than 500m form the pen. By February, when the shooting season had ended and so too much gamekeeping effort to direct and retain birds, 7% of the fixes (weighting for numbers of birds tracked and accounting for deaths over time) were within 500m from the release pen, and 10% of fixes were more than 500m.

**Figure 2.**
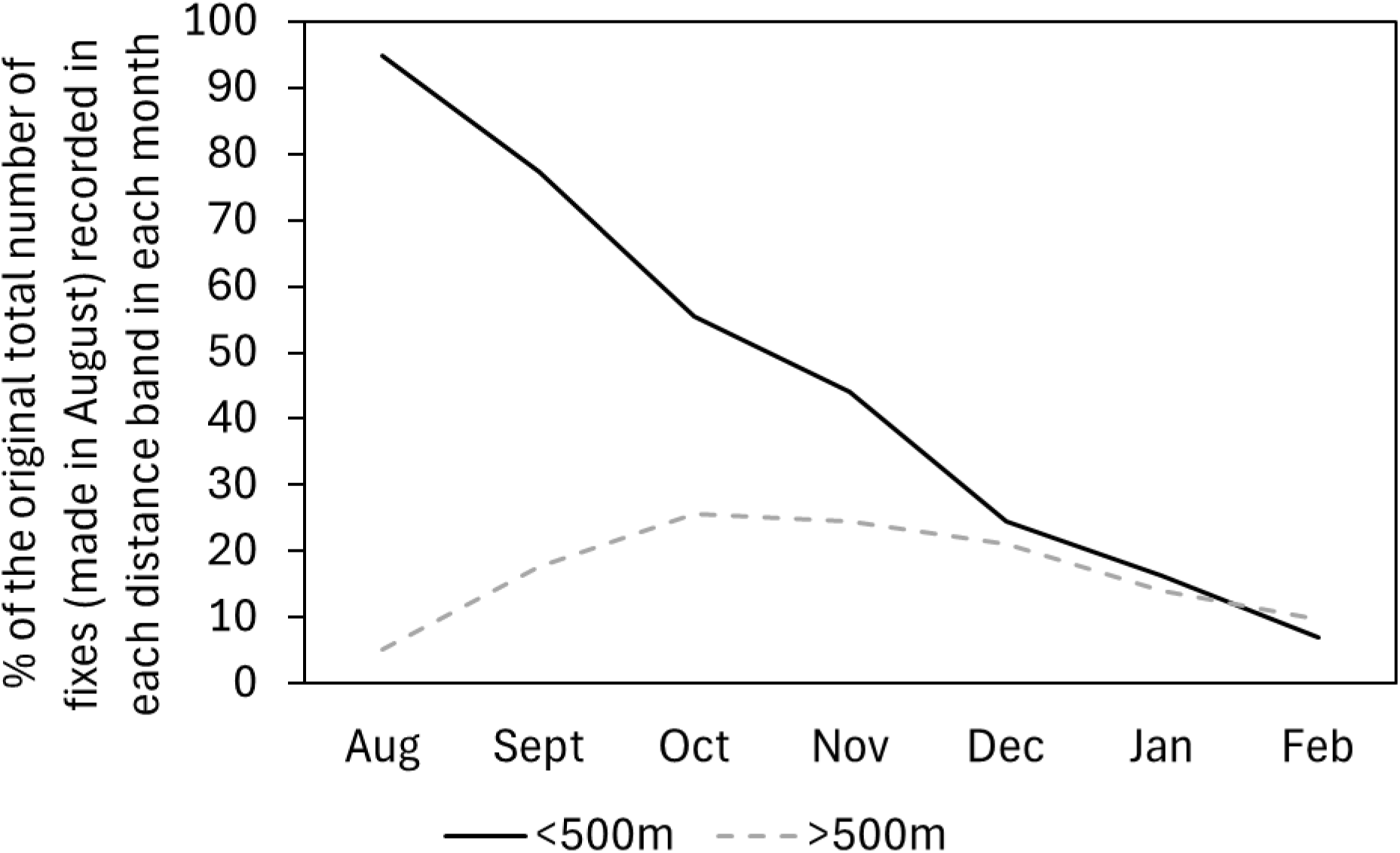
Changes in the fixes recorded from 766 released pheasants, followed in three studies, as they disperse from release pens over seven months. Fixes are shown as being recorded in an area either <500M or >500m from the release pen, to match the radius of the buffer zones currently in operation around English protected areas. In the first month, August, the percentage of fixes sum to 100%, but as the months progress, the total fixes decline as birds die, and we account for this by adjusting monthly fix totals by the original August total. Thus, in February, the total fixes are only 17% of those recorded in August, with 7% occurring in the area <500m from the release pen and 10% occurring >500m from the release pen.

We used these fix records to estimate numbers of the released birds found in protected areas at different distances in each month (Table S1, Figure 3). In August, we estimated that there might be 203,574 birds in protected areas (SSSI) within 2km of the release sites, with 186,470 birds found in protected areas <500m from release sites. If we scale up these numbers to a national level, this suggests that between 525,000 and 784,000 pheasants might be found within protected areas very soon after release. Of these, between 44,000 and 66,000 might have rapidly moved into protected areas >500m from release sites, representing 0.14% of all released pheasants. By December, the birds had dispersed further, but many had died. We now estimate 130,302 of the 9.96 million reported released birds in protected areas within 2km of the release sites, with 50,889 birds found in protected areas <500m from release sites. If we scale up these numbers to a national level, this suggests that between 336,000 and 502,000 pheasants might be found within protected areas in mid-winter. Of these, between 205,000 and 307,000 might have reached protected areas >500m from release sites, representing up to 0.8% of all released pheasants. By the end of February, a month after the shooting season and much game management has ended, we estimate that there are 50,857 of the 9.96 million reported released birds in protected areas within 2km of the release sites, with 14,817 birds found in protected areas <500m from release sites. If we scale up these numbers to a national level, this suggests that between 131,000 and 196,000 pheasants might be found within protected areas at the start of the mating season. Of these, between 93,000 and 139,000 might have reached protected areas >500m from release sites, representing up to 0.3% of all released pheasants.

**Figure 3.**
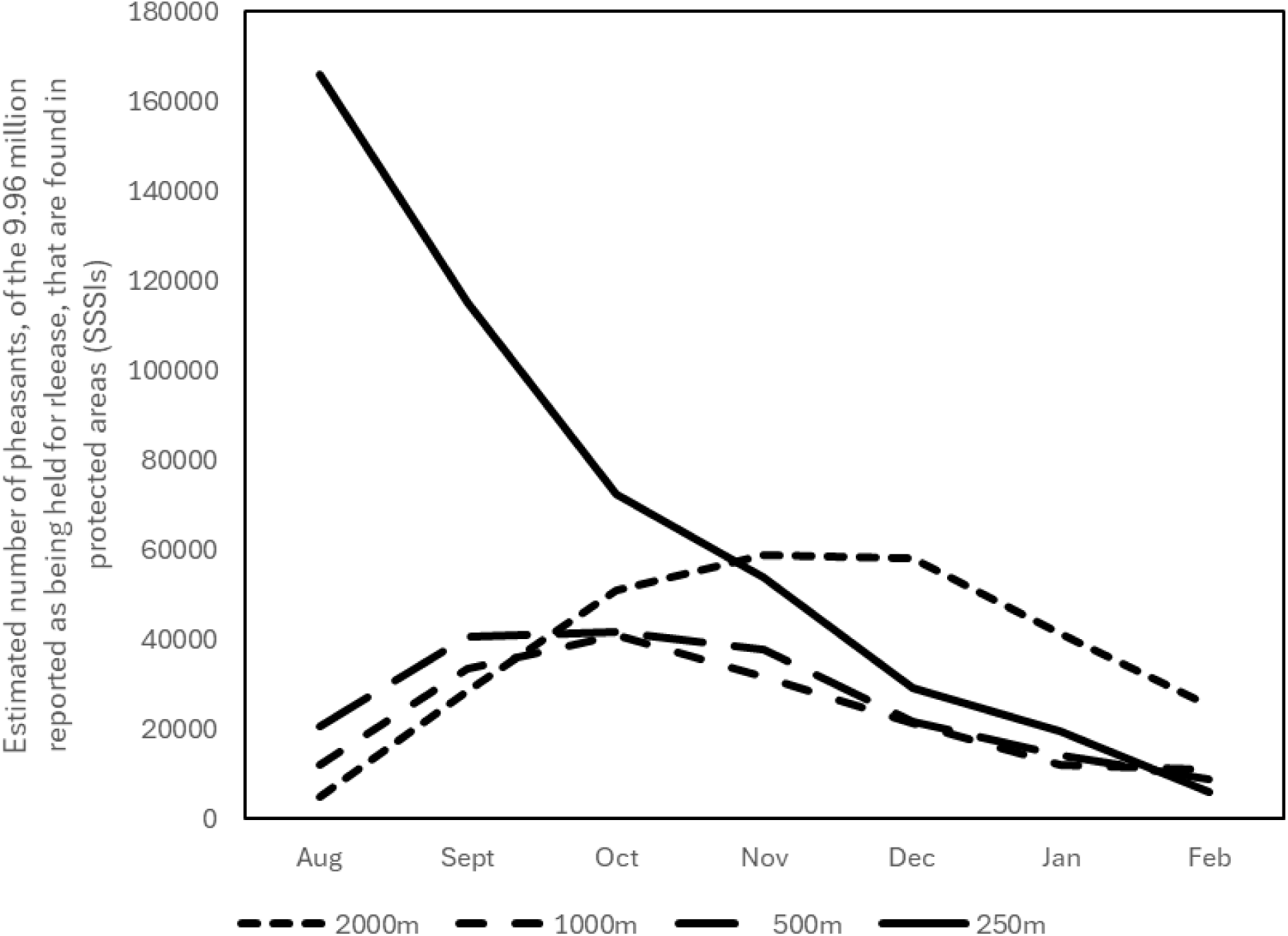
The estimated number of the 9.96 million released pheasants reported in the poultry register that might be found inside English protected areas (SSSI including SACs and SPAs) within four distance bands (black lines) of the release sites over the seven months following their release. To extrapolate these numbers to national levels, the estimates can be multiplied by 2.58 to correspond to release estimates by Madden (2021)) or 3.85 to correspond to release estimates by Aebischer (2019).

When considering estimates specific to SACs and SPAs (scaling for the total number of pheasants released), in August, there are likely to be 139,000-208,000 pheasants in SACs <2km from release sites and 168,000-251,000 in SPAs <2km from release sites. By December, estimates are 121,000-181,000 in SACs and 124,186 in SPAs. By the end of February, estimates are 49,000-73,000 in SACs and 49,000-74,000 in SPAs (Table S1, Figure S1).

Based on these estimated numbers, we calculated densities of released birds in protected areas. These varied greatly between sites, depending on whether they had protected areas nearby, how far the protected areas were from the release sites, and on the size and shape of the protected area. Overall, the maximum density estimated in any protected area (SSSI) was 2192 pheasants/ha at one site in August where the protected area was <250m from the release site. For protected areas >500m from a release site, the maximum density was 67 birds/ha at one site in October. None of the 2885 sites produced densities >70 birds/ha in PAs >500m from the release site in any month, with a maximum of 60 (2%) of the sites producing densities >7 birds/ha >500m from the release site (Table S3).

Overall, mean densities in protected areas <250m from release sites declined markedly from 12 birds/ha (SD = 87.9) in August, through 3.8 birds/ha in November to 0.4 birds/Ha in February (Figure 4). In August, the mean density of pheasants in protected areas (SSSI) >500m from release sites was 0.18 birds/ha (SD = 0.9). By December, the mean density of pheasants in protected areas (SSSI) >500m from release sites had risen to <0.32 birds/ha (SD = 1.58). By February, the mean density of pheasants in protected areas (SSSI) >500m from release sites was <0.16 birds/ha (SD = 0.81). Patterns of Natura 2000 sites (SACs and SPAs) were qualitatively similar but markedly lower, with mean August densities of <4.8 birds/ha for SACs and <2.9 birds/ha for SPAs (Figure S2). For SACs, a maximum of 26 (0.9%) sites hosted densities of >7 birds/ha >500m from the release site (in October), falling to two sites by February (Table S2). For SPAs, a maximum of 12 (0.4%) sites hosted densities of >7 birds/ha >500m from the release site (in October), falling to one site by February (Table S3).

**Figure 4.**
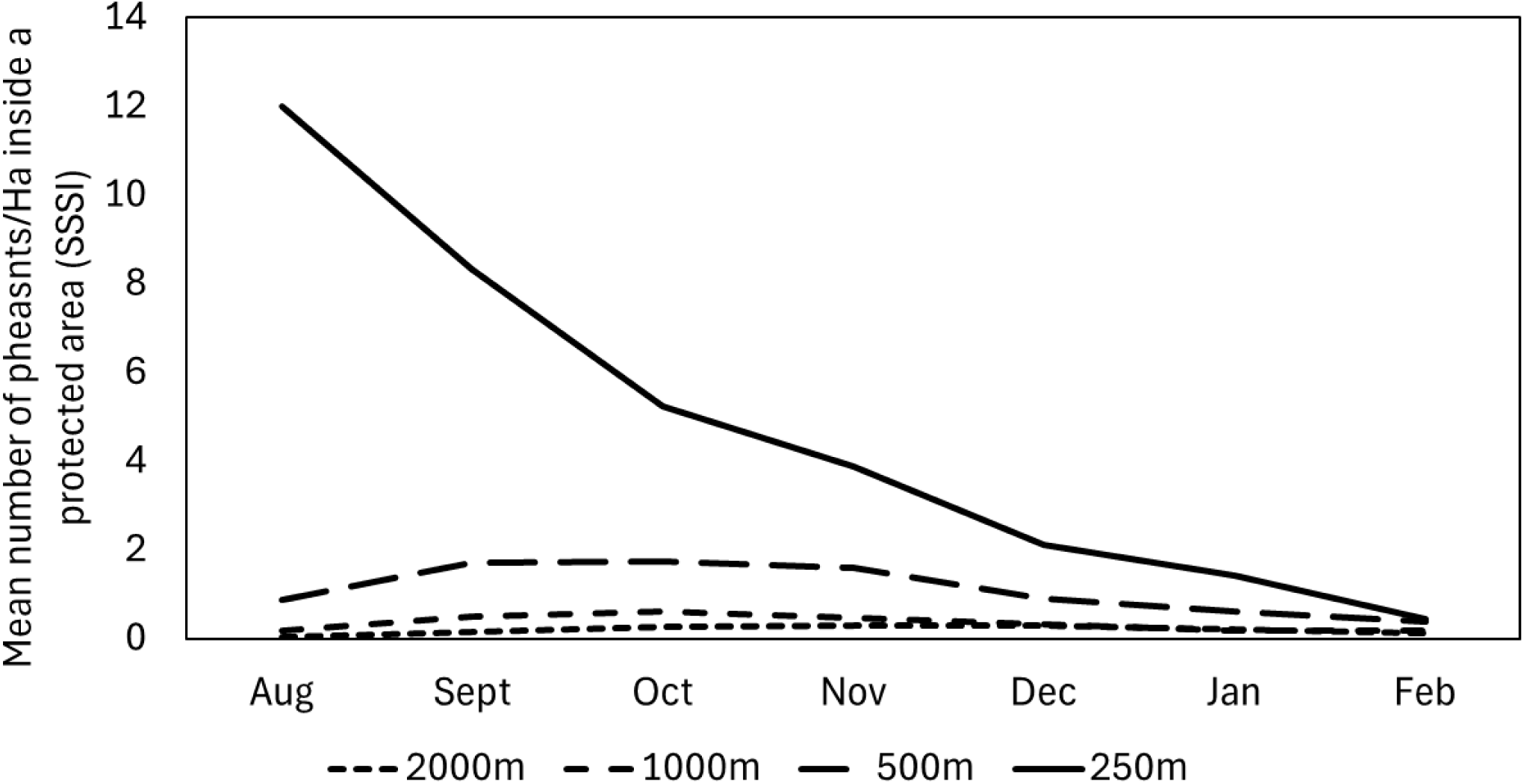
The mean densities (birds/Ha) of pheasants released at 2885 sites reported in the poultry register that might be found inside English protected areas (SSSI including SACs and SPAs) within four distance bands (black lines) of the release sites over the seven months following their release.

## DISCUSSION

Released pheasants gradually disperse from woodland pens into the surrounding countryside. However, due to high mortality soon after release, and perhaps due to supplementary feeding and resource provision in and around the release pen, their dispersal into areas >500m from the pen is relatively slow and does not exceed 26% of the original levels of fixes, falling to 10% by the end of the shooting season. By the start of the shooting season (October), 22% of the released birds were already dead. At the height of the season in December, 41% of the birds remained and by the end of February less than 15% were alive.

Based on these patterns of survival and dispersal, we derived local and national estimates of released pheasant intrusions into PAs. The strong skew in the distribution of pheasant release numbers in the UK, with a few shoots releasing a large number of birds while most shoots release relatively few, means that the effects of releases on local protected areas is likely to be highly site-specific. Further, less than 20% of shoots are located close to (<500m) PAs, involving just under 4000ha of SSSI, while just over a half of releases are within 2km of any PA, involving 118,500ha of PA. For Natura 2000 sites, this includes 59,000ha of the 546,000ha (11%) of SACs and 58,000ha of the 506,000ha (11%) of SPAs in England. Broadly speaking, a relatively small number of shoots might each release lots of pheasants close to a relatively small amount of the total protected area in England. This produces intense, but highly localised patterns of intrusion into PAs. Overall, very soon after release in August, there may be between 525,000 and 784,000 pheasants found in English protected areas including up to 168,000-251,000 in SPAs and 139,000-208,000 SACs, with the vast majority intruding into protected areas close (<500m) to release sites. Numbers intruding into protected areas >500m from release sites increase to a peak between October and December, before declining towards the end of the shooting season so that by the end of February, between 131,000 and 196,000 pheasants might remain in English protected areas, comprising ∼0.5% of all the birds originally released.

Negative ecological effects are likely to be influenced by pheasant density (e.g. Sage et al. 2005a, Gortazar et al. 2006, Pressland 2009, Neumann et al. 2015, Capstick et al. 2019, but see Davey 2008). The highest densities in protected areas occurred in August, immediately after the birds were released and had yet to disperse far or die, and in areas very close to the release sites. As the season progressed, the released birds dispersed, resulting in initial increases in densities in protected areas further (>500m) from the release sites. These densities peaked by November and then declined as the birds died. We don’t expect densities to change much after February because relatively few birds are still alive and as the mating season begins, male pheasants start to establish territories. The density estimates that we calculated correspond well with those observed by Sage *et al*. (2025) at ten UK shoots. At land managed for shooting (likely equivalent to that <500m from a release pen), they recorded densities of ∼500 birds/km^2^ soon after release, while we estimated densities of ∼640 birds /km^2^. By the end of the shooting season, they recorded densities of 100-250 birds/km^2^, while we estimated densities ∼ 80 birds/km^2^. On land not managed for shooting but within 2km of a release site and 0.5-2.5km from managed game areas, they recorded densities of <20 birds/km^2^ both around the time of release and after the end of the season, while we estimated peak densities of ∼11 bird/km^2^ at these times at >500m from release sites. At these sites during the shooting season, they recorded densities of 20-45 birds/km^2^ while we estimated densities of ∼30 birds/km^2^. This gives some confidence that the estimations that we calculated correspond reasonably well to direct measures.

Ecological damage is likely to be evident and persistent at sites where densities >700 birds/ha are present (Sage *et al*. 2021), and this forms the basis of current English release legislation. In studies that have looked at ecological effects within release pens, densities have been around two orders of magnitude higher than the mean densities (12.0 birds/ha) we calculated as occurring in PAs within 250m of release pens. Sage *et al*. (2005) reported a mean figure of 2,250 birds per hectare of pen in a 1988 sample of around 40 sites and a mean of 1,800 in a 2004 sample of 50 sites. Neumann *et al*. (2015) recorded a mean of 1,500 birds per hectare of pen at 37 sites in 2006. In our study, thirteen (0.5%) of the 2885 sites produced densities in protected areas of >700 birds/ha up to 250m away from release sites in August. This fell to six sites in September and one in October and November. Such densities were never estimated in protected areas >250m from release sites. Specific to SACs and SPAs, there were 5 sites producing densities >700 birds/ha, again, only soon after release (in August and September) and in protected areas >250m from release sites. For protected areas >500m from release sites (representing those outside the current buffer zone) mean maximum densities were 0.6 birds/ha, being three orders of magnitude lower than currently advisable release densities, and the maximum density occurring at any single site being 67 birds/ha being an order of magnitude lower than currently advisable release densities. Crucially, these maximum densities occurred over the autumn/winter, with declines as birds were shot or died naturally, such that by the start of spring, mean maximum densities >500m from release sites were all <1 bird/ha and unlikely to increase throughout the spring/summer, and the maximum density occurring at any single site being 18 birds/Ha. These estimates suggest that any density-dependent negative direct effects of released pheasants are likely to be 10-1000 times less marked than those seen within well-managed release pens.

Contrary to our understanding of the effects of high pheasant densities within release pens, we currently lack data on what ecological effect these much lower densities of pheasants might have on the environment or how long any such effects might persist. Therefore, whether the densities that we calculate might pose ecological risks is unclear. Madden et al. (2026) found mixed evidence of effects away from release pens, with reductions in seedling/sapling numbers, vascular plant richness and decayed wood up to 500m from woodland release pens, but no detectable effects on soil nutrient levels, bare ground cover and ancient woodland indicator species, all of which were previously reported to decline inside pens with high release densities. This suggests that at these distances, densities are not high enough to produce detectable negative ecological effects. However, some negative effects may not be density dependent e.g. support for predator populations may depend more on total numbers of prey available than their density if the predator covers a large home range; disease transmission may depend more on local clustering of birds than densities over larger areas. Evidence for these may come from some of the landscape scale patterns of changes in non-game fauna reported by Madden et al. (2023) associated with release sites but that are unrelated to release sizes. Greater detail of intra- and inter-specific interactions are needed to understand how these other negative effects might scale with release numbers, time of year and distances from the release pen.

Our calculations assume totally uniform dispersal from sites of release. However, pheasants exhibit strong habitat preferences (Hill & Ridley 1987, Robertson et al. 1993a,b, Lachlan & Bray 1976) and appear to follow linear features when moving (Beardsworth et al. 2021). Therefore, our simplistic approach that makes no assumptions about specific behaviours of habitats is likely to underestimate densities at some specific locations if they are especially attractive to birds e.g. feeders or cover, or because birds are concentrated in an area due to funnelling by habitat features or deliberate herding by gamekeepers. However, feeding (and other game management) within or close to PAs operates under licence or consent and one stipulation is that there is at least one feeder per 60 birds released to reduce concentrations. A fuller understanding would incorporate other aspects of pheasant or gamekeeper behaviour, coupled with detailed information about local habitat and patterns of connectivity. A modelling approach may permit generalisations to be made.

Our calculations assume that all pheasants in PAs are immigrants from gamebird releases in that year. However, there might also be survivors from previous years or wild-born offspring of released pheasants or their ancestors. Separating out released from “wild” pheasants is difficult (Jensen et al. 2012). Breeding success of released birds is generally low (Hill & Robertson 1988, Sage et al. 2003). However, wild-born pheasants survive better than released ones (Musil & Connelly 2009, Brittas et al. 1992) so once a population is established, it may persist. Without accurate data on the size and distribution of wild pheasant populations and up-to-date breeding success data, or data on movement behaviour of adult (>1year old) pheasants we cannot incorporate these additional sources of pheasants in PAs into our estimates.

As with all work in this area, our estimates are limited by the accuracy and completeness of national data on gamebird releases. We can, and do, attempt to correct for the volume of released pheasants by crude multiplication, but this cannot account for missing sites. This is critical because there is likely to be a skewed distribution such that, if large releases are conducted close to PAs, intrusion rates will be much higher. Perhaps 1/3 - 1/2 of English shoots are included in the Poultry Register (see Madden 2023b) and the omission or overrepresentation of large shoots close to PAs may change our findings. We might expect that shoots near to PAs are over-represented in the Poultry Register because currently shoots wishing to release within buffer zones or inside the PA itself may do so only under licence, and this may prompt them to also be more diligent in their compliance with Poultry Register reporting requirements. If this is the case, then we have probably overestimated the proportion of shoots close to PAs.

In conclusion, most released pheasants spend their (relatively short) lives close to the site where they were released, either because they die young and/or they don’t move very far. Consequently, most negative ecological effects attributable to them are unlikely to extend or persist very far from release sites. However, a broad-brush approach to understanding their movement, based on assumptions of uniform movement behaviour, risks missing out on local concentrations of birds which could cause local damage. To account for this, it is necessary to account for fine-scale, habitat and management dependent movement decisions and consider ecological effects that are not density dependent.

## Supporting information

ESM

## ACKNOWLEDGEMENTS

This work was funded equally by the Animal and Plant Health Agency (APHA) and the British Association for Shooting and Conservation (BASC). GPS tracking data were kindly supplied by APHA. Radio-telemetry tracking data were kindly supplied by the Game and Wildlife Conservation Trust (GWCT) and originally collected by Claire Turner. Mark Whiteside and Christine Beardsworth led the collection of ATLAS data. DEFRA provided access to the Poultry Register data under a data sharing agreement. We are very grateful to Marnie Lovejoy and Cat McMicol (BASC) and Graham Smith, Dan Warren and Dave Parrott (APHA) for advice, discussion and analytical input during the project. APHA commissioned this research as part of DEFRA’s Gamebird Research Programme.

## DATA AVAILABILITY

The data supporting this MS and the R code used in the analyses are available from https://github.com/JoahMadden/estimate-pa-release-densities. They will be placed in a permanent repository on acceptance of the paper.

