## Supplementary material for "The dispersal of released pheasants and the risk of their intrusion into English protected areas": ESM

A)

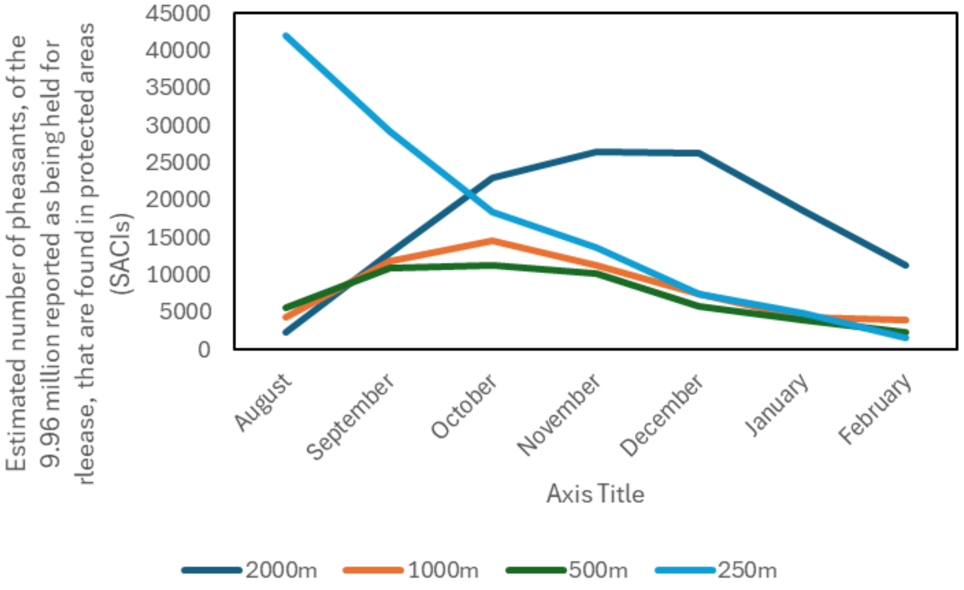


B)

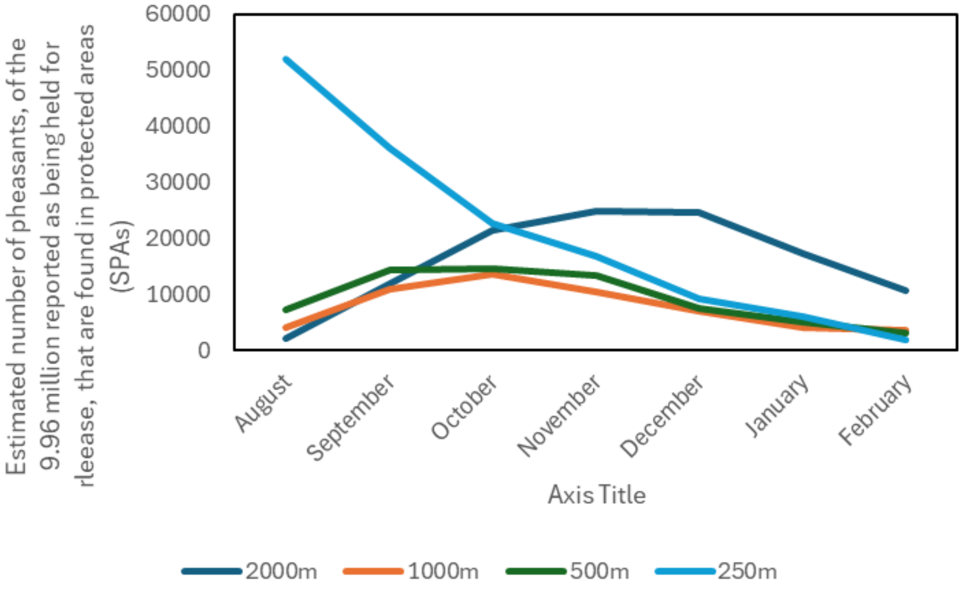


**Figure S1** The estimated number of the 9.96 million released pheasants reported in the poultry register that might be found inside English protected areas including A) SACs and B) SPAs within four distance bands (coloured lines) of the release sites over the seven months following their release. To extrapolate these numbers to national levels, the estimates can be multiplied by 2.58 to correspond to release estimates by Madden (2021)) or 3.85 to correspond to release estimates by Aebischer (2019).

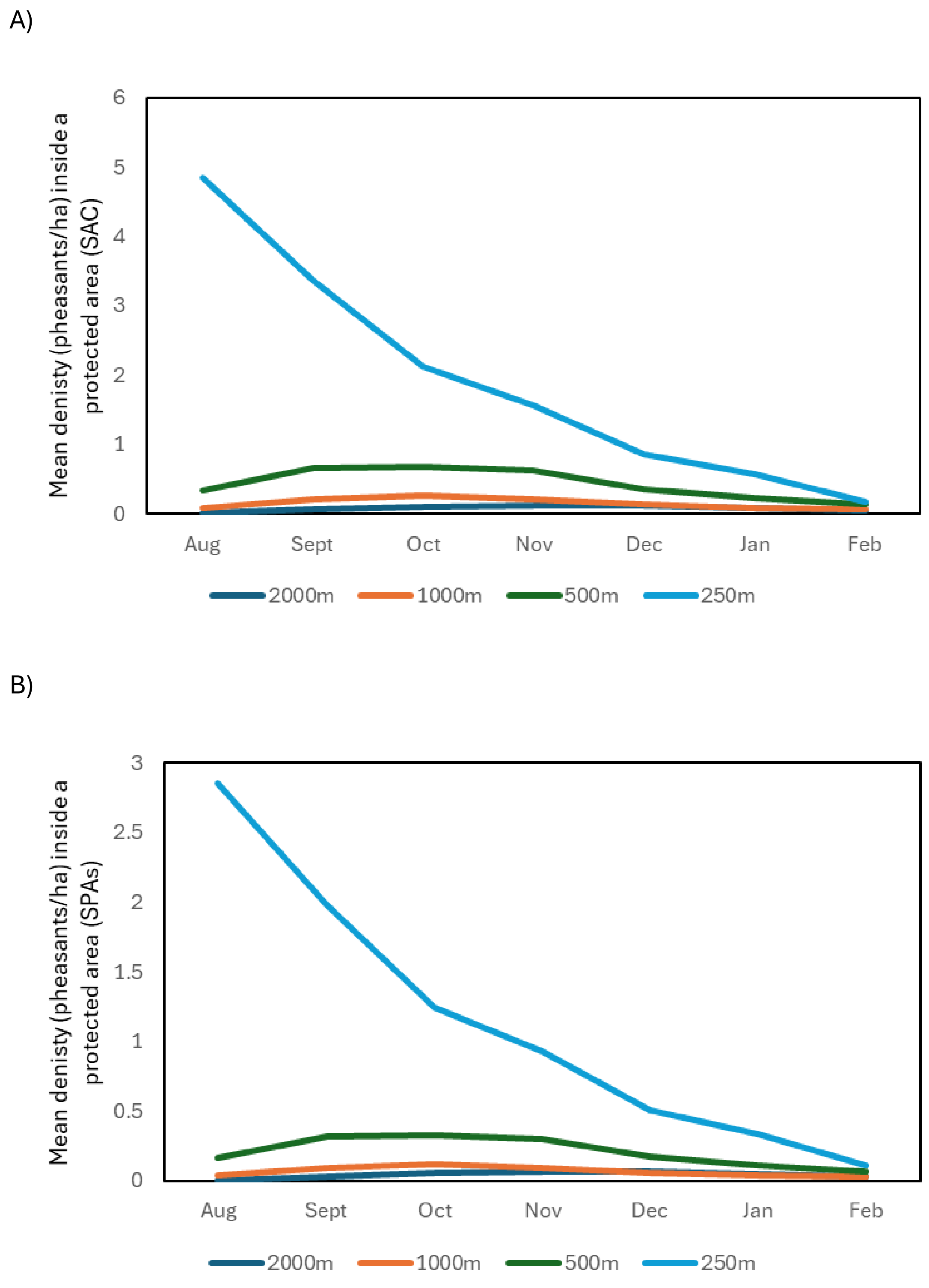


**Figure S2** The mean densities (birds/ha) of pheasants released at 2885 sites reported in the poultry register that might be found inside English protected areas including A) SACs and B) SPAs within four distance bands (coloured lines) of the release sites over the seven months following their release.

| Month | SSSI 2000m | 1000m | 500m | 250m | SAC 2000m | 1000m | 500m | 250m | SPA 2000m | 1000m | 500m | 250m |
|---|---|---|---|---|---|---|---|---|---|---|---|---|
| August | 4954 | 12149 | 20522 | 165949 | 2230 | 4283 | 5542 | 42004 | 2082 | 4003 | 7209 | 51973 |
| September | 28481 | 33322 | 40476 | 115336 | 12817 | 11747 | 10932 | 29193 | 11969 | 10978 | 14219 | 36122 |
| October | 50864 | 41088 | 41701 | 72433 | 22890 | 14485 | 11263 | 18334 | 21376 | 13536 | 14649 | 22685 |
| November | 58788 | 31839 | 37650 | 53691 | 26456 | 11224 | 10168 | 13590 | 24706 | 10489 | 13226 | 16815 |
| December | 58224 | 21188 | 21543 | 29346 | 26202 | 7470 | 5818 | 7428 | 24469 | 6980 | 7568 | 9191 |
| January | 41194 | 12145 | 14269 | 19455 | 18538 | 4282 | 3854 | 4924 | 17312 | 4001 | 5013 | 6093 |
| February | 25120 | 10923 | 8683 | 6131 | 11305 | 3851 | 2345 | 1552 | 10557 | 3598 | 3050 | 1920 |

**Table S1** The estimated number of the 9.96 million released pheasants reported in the poultry register that might be found inside English protected areas (SSSI, SAC, SPA) within four distance bands of the release sites over the seven months following their release. To extrapolate these numbers to national levels, the estimates can be multiplied by 2.58 to correspond to release estimates by Madden (2021)) or 3.85 to correspond to release estimates by Aebischer (2019).

| Month | SSSI 2000m | 1000m | 500m | 250m | SAC 2000m | 1000m | 500m | 250m | SPA 2000m | 1000m | 500m | 250m |
|---|---|---|---|---|---|---|---|---|---|---|---|---|
| August | 0.03 | 0.18 | 0.86 | 11.99 | 0.01 | 0.08 | 0.34 | 4.84 | 0.01 | 0.03 | 0.16 | 2.86 |
| September | 0.15 | 0.50 | 1.70 | 8.33 | 0.06 | 0.21 | 0.66 | 3.37 | 0.03 | 0.09 | 0.32 | 1.98 |
| October | 0.26 | 0.61 | 1.75 | 5.23 | 0.10 | 0.26 | 0.68 | 2.11 | 0.05 | 0.11 | 0.33 | 1.25 |
| November | 0.30 | 0.47 | 1.58 | 3.88 | 0.12 | 0.20 | 0.62 | 1.57 | 0.06 | 0.09 | 0.29 | 0.92 |
| December | 0.30 | 0.32 | 0.90 | 2.12 | 0.11 | 0.13 | 0.35 | 0.86 | 0.06 | 0.06 | 0.17 | 0.50 |
| January | 0.21 | 0.18 | 0.60 | 1.41 | 0.08 | 0.08 | 0.23 | 0.57 | 0.04 | 0.03 | 0.11 | 0.33 |
| February | 0.13 | 0.16 | 0.36 | 0.44 | 0.05 | 0.07 | 0.14 | 0.18 | 0.03 | 0.03 | 0.07 | 0.11 |

**Table S2** The estimated mean densities (birds/ha) of pheasants released at 2885 sites reported in the poultry register that might be found inside English protected areas (SSSI, SAC, SPA) within four distance bands of the release sites over the seven months following their release.

| | | Sites with >700 birds/Ha | | | | Sites with >70 birds/Ha | | | | Sites with >7 birds/Ha | | | |
|---|---|---|---|---|---|---|---|---|---|---|---|---|---|
| | | 2000m | 1000m | 500m | 250m | 2000m | 1000m | 500m | 250m | 2000m | 1000m | 500m | 250m |
| **SSSI** | August | 0 | 0 | 0 | 13 | 0 | 0 | 7 | 86 | 0 | 11 | 78 | 198 |
| | September | 0 | 0 | 0 | 6 | 0 | 0 | 12 | 71 | 3 | 49 | 120 | 187 |
| | October | 0 | 0 | 0 | 1 | 0 | 0 | 13 | 56 | 8 | 60 | 120 | 175 |
| | November | 0 | 0 | 0 | 1 | 0 | 0 | 10 | 48 | 19 | 43 | 106 | 172 |
| | December | 0 | 0 | 0 | 0 | 0 | 0 | 7 | 26 | 19 | 27 | 82 | 129 |
| | January | 0 | 0 | 0 | 0 | 0 | 0 | 2 | 14 | 5 | 11 | 55 | 100 |
| | February | 0 | 0 | 0 | 0 | 0 | 0 | 2 | 1 | 3 | 9 | 31 | 48 |
| **SAC** | August | 0 | 0 | 0 | 5 | 0 | 0 | 1 | 43 | 0 | 4 | 34 | 88 |
| | September | 0 | 0 | 0 | 1 | 0 | 0 | 2 | 33 | 1 | 22 | 57 | 84 |
| | October | 0 | 0 | 0 | 0 | 0 | 0 | 2 | 25 | 3 | 26 | 57 | 80 |
| | November | 0 | 0 | 0 | 0 | 0 | 0 | 2 | 21 | 5 | 18 | 49 | 76 |
| | December | 0 | 0 | 0 | 0 | 0 | 0 | 1 | 9 | 5 | 8 | 36 | 57 |
| | January | 0 | 0 | 0 | 0 | 0 | 0 | 1 | 6 | 2 | 4 | 23 | 48 |
| | February | 0 | 0 | 0 | 0 | 0 | 0 | 1 | 0 | 1 | 2 | 12 | 21 |
| **SPA** | August | 0 | 0 | 0 | 5 | 0 | 0 | 1 | 22 | 0 | 2 | 15 | 46 |
| | September | 0 | 0 | 0 | 1 | 0 | 0 | 2 | 16 | 1 | 10 | 27 | 45 |
| | October | 0 | 0 | 0 | 0 | 0 | 0 | 2 | 13 | 3 | 12 | 27 | 43 |
| | November | 0 | 0 | 0 | 0 | 0 | 0 | 1 | 10 | 5 | 8 | 23 | 42 |
| | December | 0 | 0 | 0 | 0 | 0 | 0 | 1 | 7 | 5 | 3 | 16 | 29 |
| | January | 0 | 0 | 0 | 0 | 0 | 0 | 0 | 5 | 1 | 2 | 11 | 25 |
| | February | 0 | 0 | 0 | 0 | 0 | 0 | 0 | 0 | 1 | 1 | 6 | 10 |

**Table S3** The number of the 2885 sites reported in the poultry register where the estimated mean densities (birds/ha) of pheasants exceed 700 birds/ha (the density that is known to cause negative direct ecological effects in release pens) and two orders of magnitude below that (70 & 7 birds/ha), that might be found inside English protected areas (SSSI, SAC, SPA) within four distance bands of the release sites over the seven months following their release. For reference, the 198 SSSI sites where densities exceed 7 birds/ha in PAs within 250m of the release site represent 7% of all reported releases.
